# OMSV enables accurate and comprehensive identification of large structural variations from nanochannel-based single-molecule optical maps

**DOI:** 10.1101/143040

**Authors:** Le Li, Tsz-Piu Kwok, Alden King-Yung Leung, Yvonne Y. Y. Lai, Iris K. Pang, Grace Tin-Yun Chung, Angel C. Y. Mak, Annie Poon, Catherine Chu, Menglu Li, Jacob J. K. Wu, Ernest T. Lam, Han Cao, Chin Lin, Justin Sibert, Siu-Ming Yiu, Ming Xiao, Kwok-Wai Lo, Pui-Yan Kwok, Ting-Fung Chan, Kevin Y. Yip

## Abstract

Human genomes contain structural variations (SVs) that are associated with various phenotypic variations and diseases. SV detection by sequencing is incomplete due to limited read length. Nanochannel-based optical mapping (OM) allows direct observation of SVs up to hundreds of kilo-bases in size on individual DNA molecules, making it a promising alternative technology for identifying large SVs. SV detection from optical maps is non-trivial due to complex types of error present in OM data, and no existing methods can simultaneously handle all these complex errors and the wide spectrum of SV types. Here we present a novel method, OMSV, for accurate and comprehensive identification of SVs from optical maps. OMSV detects both homozygous and heterozygous SVs, SVs of various types and sizes, and SVs with and without creating/destroying restriction sites. In an extensive series of tests based on real and simulated data, OMSV achieved both high sensitivity and specificity, with clear performance gains over the latest existing method. Applying OMSV to a human cell line, we identified hundreds of SVs >2kbp, with 65% of them missed by sequencing-based callers. Independent experimental validations confirmed the high accuracy of these SVs. We also demonstrate how OMSV can incorporate sequencing data to determine precise SV break points and novel sequences in the SVs not contained in the reference. We provide OMSV as open-source software to facilitate systematic studies of large SVs.

## Introduction

Structural variations (SVs), defined as genomic alterations involving segments larger than 1kbp (Feuk et al., 2006), are prevalent in human genomes. They represent characteristic differences among human populations (Sudmant et al., 2015), and are associated with various diseases (Stankiewicz and Lupski, 2010; Weischenfeldt et al., 2013).

Current sequencing technologies, including second-generation and commercial third-generation sequencing platforms, produce sequencing reads from a hundred to tens of thousands of base pairs only, making it challenging to study long repetitive regions and complex structural rearrangements. For instance, some large insertions cannot be contained in a single read, and their detection requires either sequence assembly (Li et al., 2011) or reference alignment (Alkan et al., 2011; Medvedev et al., 2009), with the help of paired-end or mate-pair sequencing with large insert sizes (English et al., 2015; Zeitouni et al., 2010). In general, these methods are not ideal for detecting large SVs accurately and comprehensively, especially SVs that involve long DNA sequences not present in the reference sequence (Gudbjartsson et al., 2015; Mohiyuddin et al., 2015).

Optical mapping (OM) (Dimalanta et al., 2004) is a promising alternative technology that provides structural information of individual long DNA molecules. In nanochannel-based optical mapping (Lam et al., 2012; Seo et al., 2016), DNA molecules are digested by a nicking endonuclease to create single-strand nicks, which are then repaired with fluorescent dye conjugated nucleotides. The resulting DNA molecules are linearized in nanochannels and imaged using high-resolution fluorescent microscopy (Figure S1). The final outputs are the optical maps, which record restriction site label locations on each DNA molecule. SVs can be identified by comparing the observed label pattern with the expected pattern based on the reference sequence (Figure 1a). For example, two sites significantly farther apart on an optical map than their corresponding locations on the reference could indicate an insertion.

**Figure 1:**
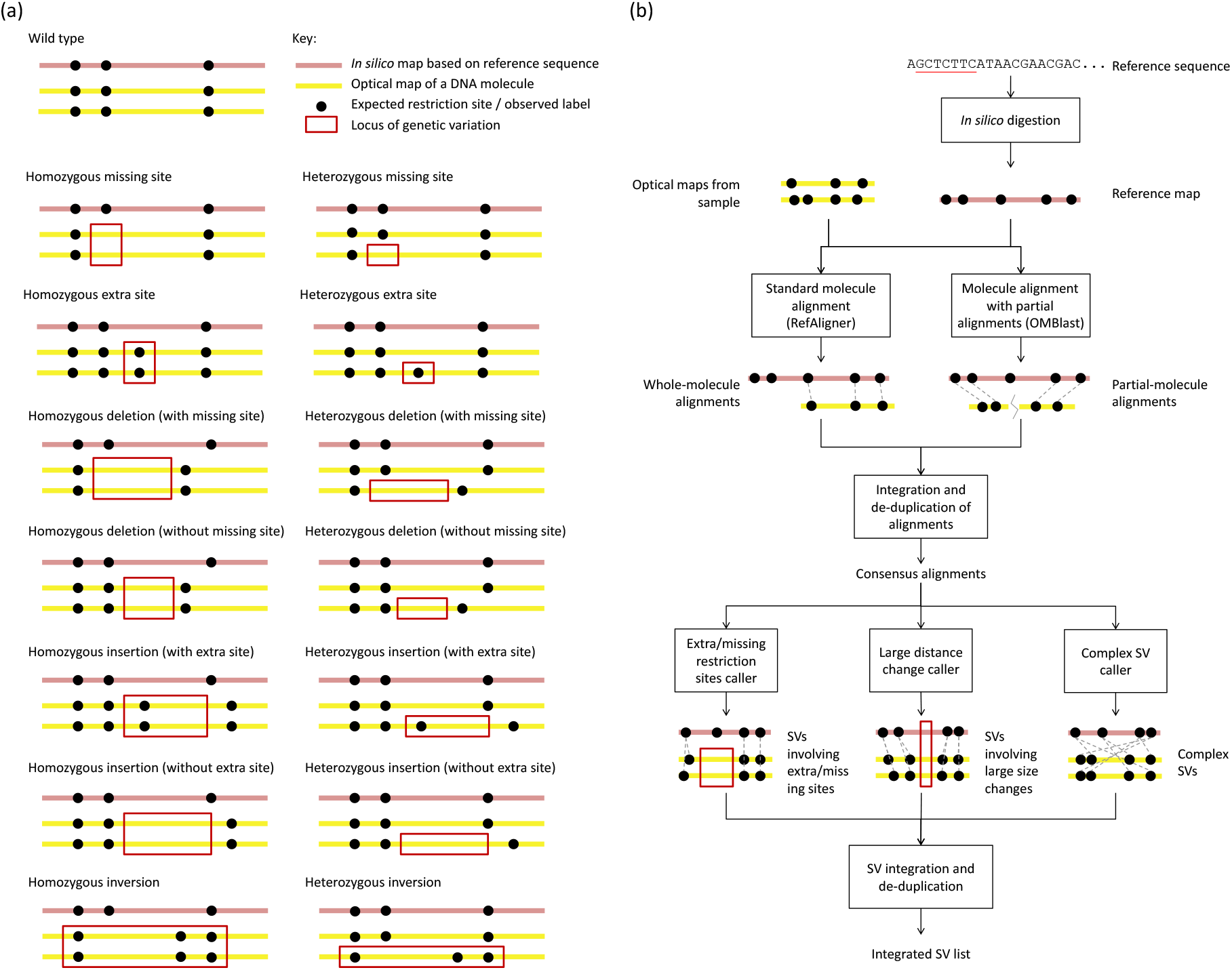
Underlying concepts of OMSV. (a) Different types of SVs and their idealized appearance patterns on optical maps. Real OM data contain various types of errors that make these patterns less apparent. Inversions are shown as an example type of complex SVs, while OMSV can also detect translocations and copy number variations. (b) The overall OMSV pipeline for identifying SVs from optical maps. Optical maps from a study sample are aligned to the reference map using two different aligners. Their results are integrated to form a single list of consensus alignments, which are then passed to three SV calling modules to identify different types of SVs.

**Figure 2:**
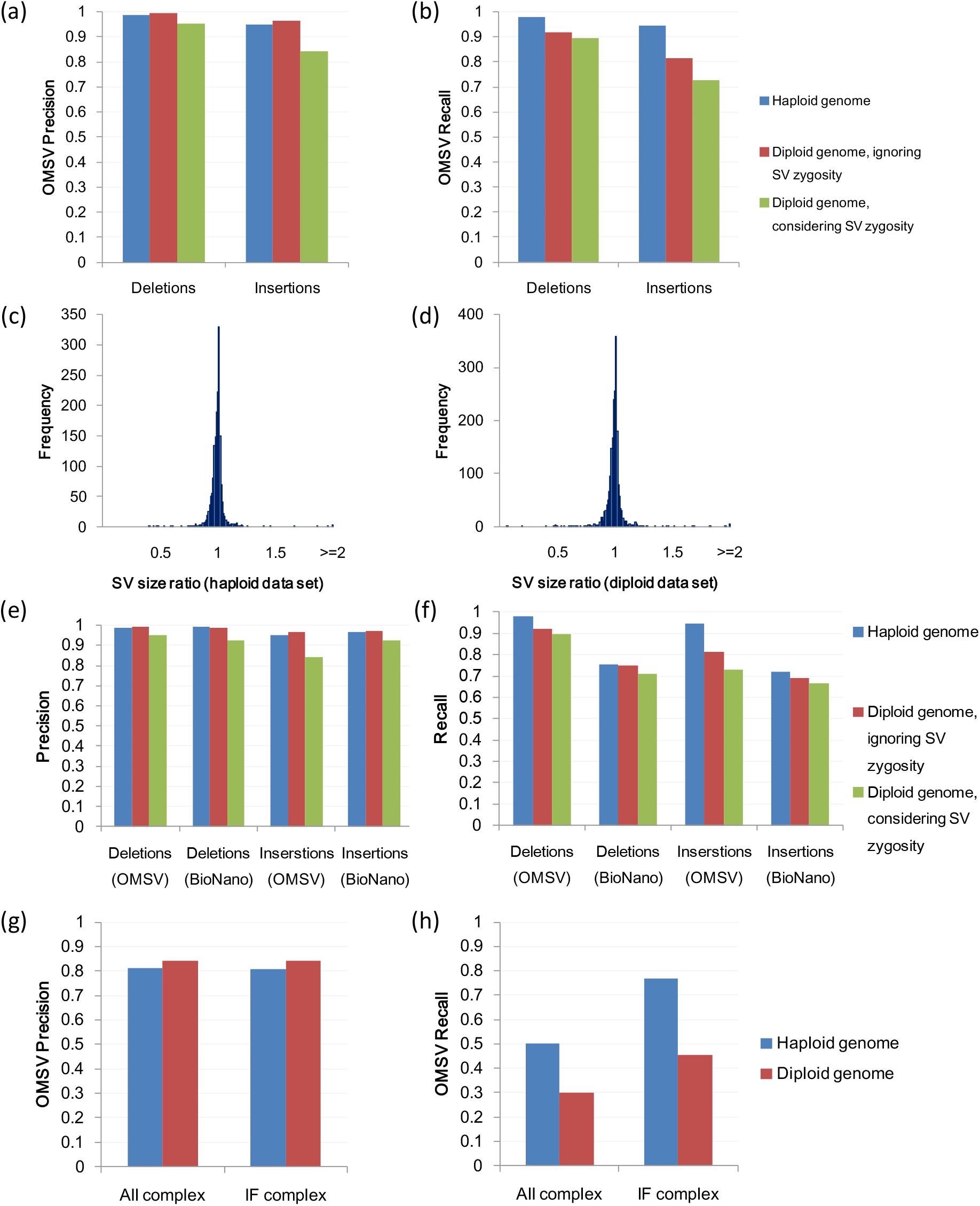
Results based on the default simulated data sets. Precision (a) and recall (b) of OMSV. Ratio of SV sizes determined by OMSV to their actual sizes, for the haploid (c) and diploid (d) data sets. Precision (e) and recall (f) of OMSV as compared to BioNano Solve. Results in Panels (a) to (f) are all based on insertions and deletions larger than 2kbp. Precision (g) and recall (h) of OMSV in calling complex SVs from the simulated data, including the whole set (“All”) and only the intrinsically feasible (“IF”) ones.

Due to the much longer length of optical maps (up to 1Mbp) compared to sequencing reads, OM has been found very powerful in SV discovery (Cao et al., 2014; Dong et al., 2013; Lam et al., 2012; Ray et al., 2013; Teague et al., 2009). Current high-throughput OM methods can produce optical maps for a hundred thousand molecules within a few hours, at an average size of several hundred kbps per molecule. These molecules can be full-length DNA derived from species with a small genome, or fragments of very long DNA molecules such as human chromosomes.

Analyzing optical maps is non-trivial due to various types of error present in the data (Tong, 2010; Valouev et al., 2006). False positives (false labels observed but not coming from true restriction sites) can occur due to non-specific enzymatic cuts or DNA breakage. False negatives (true restriction sites not observed on the optical maps) can occur due to incomplete enzyme digestion. Sizing errors (deviation between measured and actual distance between two restriction sites on an optical map) can occur due to DNA fragments that are over-stretched or not completely linearized. Finally, labels of close restriction sites may merge into a single label in the observed data due to limitations in imaging resolution. As a result of all these error types, specialized methods have been proposed for various computational tasks related to the analysis of optical maps, including error modeling (Tong et al., 2007; Tong, 2010), molecule alignment (Leung et al., 2017b; Nagarajan et al., 2008; Shelton et al., 2015; Teague et al., 2009; Valouev et al., 2006), *de novo* and reference-assisted assembly (Kim et al., 2013; Lin et al., 2012; Valouev et al., 2006), and detection of SVs (Hastie et al., 2013; Lam et al., 2012; Ray et al., 2013; Teague et al., 2009).

Existing methods for calling SVs from optical maps have several major limitations. First, most of them require a *de novo* assembly of the optical maps or the construction of a consensus map (Cao et al., 2014; Lam et al., 2012; Teague et al., 2009), making the accuracy of SV calls dependent on the reliability of these difficult procedures. Second, none of the current methods can simultaneously 1) detect both homozygous and heterozygous SVs, 2) handle SVs of a wide range of sizes, and 3) evaluate SV probabilities based on a formal error model and the optical maps that support/do not support the SVs. Besides, almost none of the existing methods have made their software publicly available, which hampers the widespread use of optical mapping in studying SVs.

Here we describe a comprehensive SV calling pipeline and corresponding open-source software, OMSV (available at the supplementary Web site, http://yiplab.cse.cuhk.edu.hk/omsv/), which overcomes these limitations. We demonstrate the effectiveness of OMSV using both simulations and optical maps produced from a family trio. In addition, we show that when OMSV was applied to detect SVs in a human cell line, many of our detected SVs were missed by typical sequencing-based SV callers. Some of our detected SVs were experimentally tested using DNA isolated from the cell line, and all of them were successfully validated. Finally, we describe how OMSV can combine optical maps and sequencing data to identify precise SV break points and uncover novel sequences involved in the SVs.

## Results

### The OMSV pipeline

OMSV contains two main steps (Figure 1b, Materials and Methods). In the first step, it aligns optical maps to the reference map, which is deduced from the reference sequence and the recognition motif of the nicking enzyme by *in silico* digestion. Two different aligners are used, including RefAligner (Shelton et al., 2015), which can efficiently align optical maps highly similar to the reference, and OMBlast (Leung et al., 2017b), which can handle more complex genomic rearrangements by split-aligning a single optical map to multiple regions on the reference. The alignment results from the two aligners are integrated to form a single set of consensus alignments. In the second step of OMSV, these alignments are passed to three separate SV calling modules for three corresponding types of SVs, namely 1) SVs involving the creation or removal of restriction sites, 2) SVs involving large distance changes between restriction sites, and 3) more complex SVs such as inversions and translocations. SVs identified from these modules are then integrated and de-duplicated to form a final list of SVs.

In the SV calling modules, a formal error model is used to compare the likelihoods of the reference genotype (i.e., no SVs), homozygous SVs, and heterozygous SVs. An SV is called only if a set of stringent criteria are satisfied (Materials and Methods).

### Simulations confirm the effectiveness of OMSV

To test the effectiveness of OMSV, we generated simulated OM data from artificial haploid and diploid human genomes, by introducing various types of genetic variants to the reference genome hg38 followed by simulation of noisy optical maps with all types of error (Materials and Methods). We defined a default error setting, and additional settings that covered a wide range of false positive and false negative rates of nicking sites and depths of coverage, leading to a total of 28 sets of simulated OM data (Tables S1,S2). The default diploid set has data properties similar to those observed in real OM data (Table S4).

Next, we applied OMSV to identify SVs from these simulated OM data sets, and compared the results to the actual lists of synthesized SVs in order to determine OMSV’s precision (fraction of called SVs that are correct) and recall (fraction of simulated SVs correctly called by OMSV).

Here we first focus on insertions and deletions (indels) larger than 2kbp in the data sets with the default setting, since they constitute a large fraction of our simulated SVs and these large SVs are difficult for short-read based methods to identify accurately. In the case of the haploid genome, both the precision and recall of OMSV were 98% for deletions and 95% for insertions (Figures 2a,b), showing that it was highly effective. In the case of the diploid genome, when the goal was to identify SV locations only without considering correctness of zygosity, OMSV again achieved high precision (99%) and recall (92%) for deletions, and high precision (97%) and moderate recall (81%) for insertions due to fewer optical maps supporting the SVs in the heterozygous cases. When correct zygosity was also required for an SV to be considered correctly called, OMSV still achieved 90% precision and 81% recall on average. For the SVs correctly identified from the two data sets, we further compared their estimated sizes by OMSV with the actual sizes up to the closest defining nicking sites, and found them to be very similar in most cases (Figures 2c,d), with a median size ratio of 1.0028 and 1.0029 for the haploid and diploid data sets, respectively.

To benchmark the performance of OMSV, we compared it with the latest version of the assembly-based SV caller BioNano Solve, which is the only other SV caller for nanochannel-based optical mapping with publicly available software. We found that the precision of the two methods was comparable, but OMSV had 10%-31% higher recall (Figures 2e,f). Moreover, since BioNano Solve required a *de novo* assembly of the optical maps, its running time was 16-21 times longer than OMSV (including the alignment time).

To evaluate the robustness of OMSV, we performed three additional sets of tests. First, using the diploid data set with default settings, we checked the SVs with various numbers of optical maps aligned to their loci and having different likelihood ratios as computed by OMSV. We found that OMSV’s precision remained highly stable at different values of these variables (Figure S2), and the default parameter values of OMSV (at least 10 aligned optical maps and a null-to-alternative likelihood ratio of at most 10^−6^ for an SV to be called) provided a good tradeoff between precision and recall. Second, we compared the performance of OMSV on data sets with different depths of coverage. To separate the effects of alignment errors and SV calling errors, we also considered an idealized situation with no alignment errors (Materials and Methods). The depth of coverage was found to have virtually no effects on the precision of OMSV for the depth values considered (Figure S3a), but it correlated with the recall (Figure S3b). Importantly, by comparing our results with and without involving optical map alignments, we found that the decreased recall at low depth of coverage was almost completely due to alignment errors but not SV calling errors. Third, we altered the false positive and false negative rates of the OM data, and found that the performance of OMSV remained stable for most settings until the error rates reached unrealistically large values not typically seen in real data (Figures S4,S5). Again, we found that the performance drop at high false positive and false negative rates correlated strongly with alignment errors, and thus the performance of OMSV should be automatically improved with better alignment accuracy. Overall, these three sets of tests show that OMSV is generally robust against different data properties and parameter settings.

We also compared different alignment strategies involving alignments from only one of the two aligners, their intersection, and their union. The results (Figure S6) show that taking the union of the two aligners had the best tradeoff between precision and recall, especially when the data set had a low depth of coverage.

For complex SVs (Figures 2g,h), OMSV achieved 80-85% precision but only 30-50% recall on the two default sets. Many of the missed SVs were found to be intrinsically infeasible to call, including inversions that contain no nicking sites or symmetric nicking site patterns that would not change upon the inversions. After filtering these cases, the recall rate of the resulting intrinsically feasible (IF) complex SVs was substantially improved to 45-80%. BioNano Solve contained a function for calling complex SVs, but failed to detect any of them from the simulated data.

Taken together, the simulation results show that OMSV can identify large SVs accurately and comprehensively on data sets with properties typical in real data.

In terms of running time of OMSV, the main bottleneck was optical map alignments (Table S3). This limitation can be overcome by running the aligners on multiple threads in parallel, leading to an overall running time of OMSV of less than 5 hours for each simulated human sample with a 100x genome coverage.

### OMSV identifies SVs concordantly from different members of a family

We next tested OMSV on the optical maps produced from a family trio in a former study (Mak et al., 2016) (Table S4). Genetic variants from this trio were previously reported (MacDonald et al., 2014), but they are mostly small variants. OMSV called 920–990 large indels from the three samples independently (Table S5, Supplementary File S1, Materials and Methods), with an average size of 6.3kbp and a maximum of 89kbp. Among these indels, 22 contain an insertion allele and a deletion allele at the same SV locus. OMSV also called 86–158 complex SVs from this trio. Since the actual SVs in these individuals were not known, we used four different methods to estimate the accuracy of OMSV.

First, we hid the sex of the samples from OMSV, and checked the number of SV calling errors related to the sex chromosomes (Table S5). When pseudo-autosomal regions were excluded, in the female samples NA12878 and NA12892, 52 and 54 SVs were called on the X chromosome, respectively, whereas no SVs were wrongly called on the Y chromosome. In terms of zygosity, the male sample NA12891 had 15 SVs wrongly called as heterozygous among the 51 SVs called on the non-pseudo-autosomal regions of the sex chromosomes. Based on these numbers, the estimated zygosity precision was (51-15)/51 = 71%.

Second, we compared the SVs called from the three individuals. Among the high-confidence calls (Materials and Methods), 98% were concordant with Mendelian inheritance when zygosity of the SVs was ignored, and 85% were concordant when zygosity was considered. We used the precision and recall values from our simulations to estimate the expected Mendelian concordance to be 96% when zygosity was ignored and 83% when zygosity was considered (Materials and Methods), suggesting that the accuracy of OMSV on the trio data was comparable to that on the simulated data.

Third, we compared our SV calls with the manual checking results from Mak et al. (2016) based on nicking site patterns of aligned molecules (Table S6). Among our SVs with manual checking results, 96-97% of them were considered correct when zygosity was ignored, which is similar to the precision values in the simulation study. When zygosity was considered, 73-74% of our SVs were considered correct by the manual checking results, which is lower than that in the simulation. Together, these results suggest that OMSV could identify SV locations accurately but determining the correct zygosity of the SVs could be more difficult with real data.

Finally, we compared our indel list from NA12878 with two lists of indels detected from this sample previously using sequencing-based methods (Parikh et al., 2016; Sudmant et al., 2015). Focusing on large (>2kbp) indels, the intersection of OMSV’s list and either of these two sequencing-based lists (81 and 90 indels, respectively) was similar to the intersection of these two lists (84 indels) (Figure S7). Interestingly, 500 (96%) of the insertions and 178 (38%) of the deletions called by OMSV were unique. Based on the estimated accuracy of OMSV, many of these novel indels are expected to be real. These observations suggest that OMSV is able to identify SVs commonly called by other sequencing-based methods as well as uncover novel ones missed by them.

We select two examples to illustrate the SVs identified by OMSV. In the first example on chromosome 6 (Figure 3a), the father (NA12891) has a heterozygous insertion of around 14.6kbp, the mother (NA12892) has a heterozygous insertion of around 22.7kbp, and the daughter (NA12878) inherits both insertions from the parents. This example demonstrates the abilities of OMSV in identifying heterozygous SVs and loci with two distinct alleles both different from the reference. In the second example (Figure 3b), a large inversion of around 123.3kbp was consistently found on chromosome X from all three individuals, with clear nicking site patterns that support the inversion.

**Figure 3:**
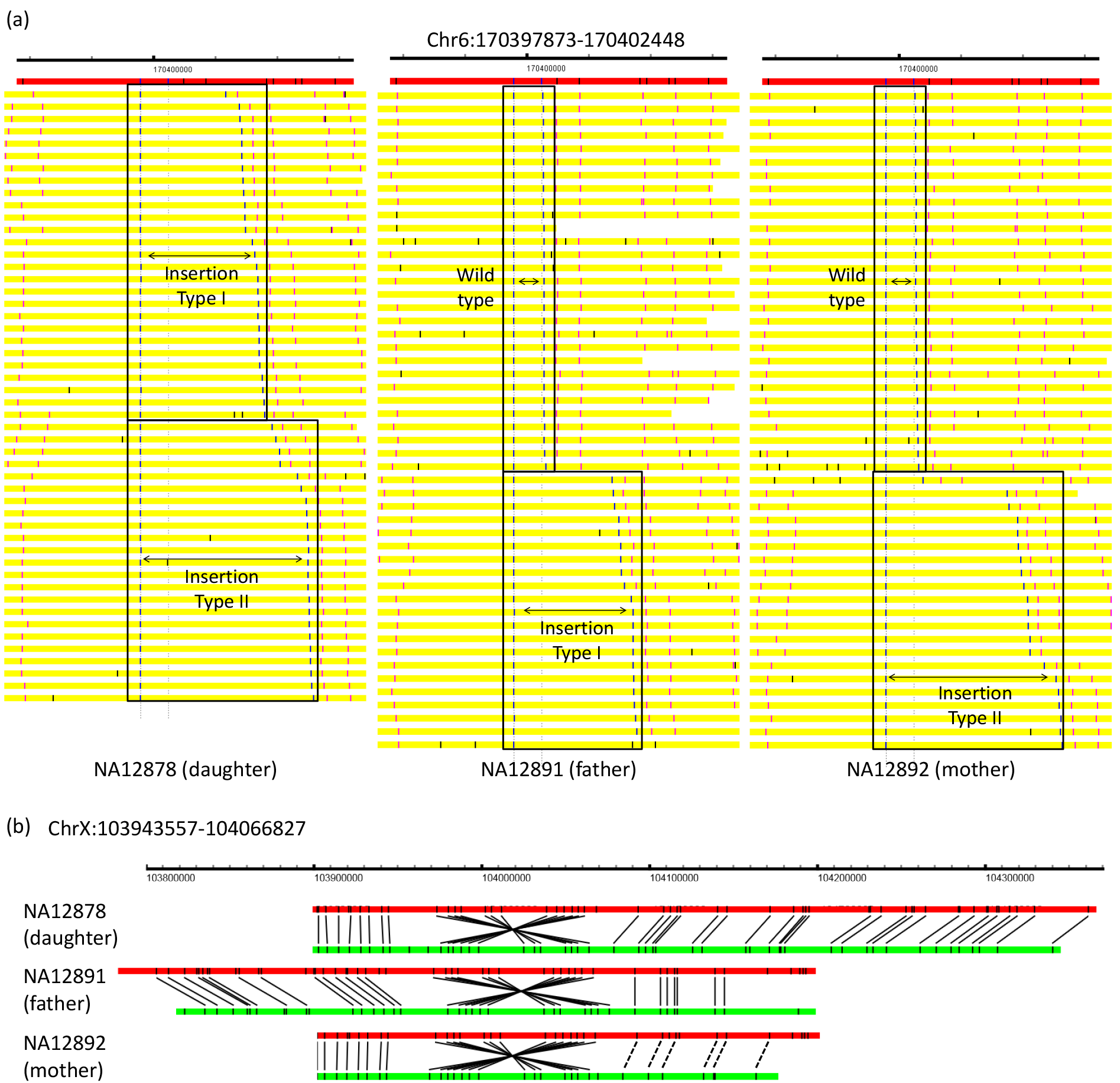
Examples of SVs identified from the trio. (a) An insertion sites identified on chromosome 6, visualized by OMView (Leung et al., 2017a) using the anchor view with the nicking site immediately before the insertion served as the anchor. The red horizontal bars show the reference, with the nicking sites marked in black vertical lines. Each yellow horizontal bar represents an optical map, with the two aligned nicking site labels defining the SVs in blue, other aligned labels in pink, and unaligned labels in black. For each individual, optical maps are arranged into different sections based on the allele that they support. The father has a heterozygous insertion of around 14.6kbp (“Insertion Type I”). The mother has a heterozygous insertion of around 22.7kbp (“Insertion Type II”). The daughter inherited both insertions from her parents. (b) An inversion identified on chromosome X, visualized using the alignment view of OMView. For each individual, the top horizontal bar shows the reference and the bottom horizontal bar shows a representative optical map. Black solid and dashed lines linking the reference and the optical map respectively represent aligned nicking sites and nicking sites that should probably be aligned but missed by the alignment pipeline.

### OMSV identifies many SVs missed by short-read based SV callers

To further evaluate the ability of OMSV in detecting novel SVs, we produced optical maps from the human C666-1 cell line (Cheung et al., 1999) (Table S7). C666-1 cells consistently harbor multiple Epstein Barr virus (EBV) episomes. As a first check of the data produced, we aligned the optical maps to the EBV reference in C666-1 (Tso et al., 2013), and found a large number of well-aligned optical maps (Figure S8). Comparing the average depth of coverage of the optical maps aligned to the human (72x) and EBV (847x) references, we estimated an average of 24 copies of the EBV genome per C666-1 cell, which is highly consistent with a previous estimate based on sequencing data (Xiao et al., 2016).

We then applied OMSV to identify SVs in the C666-1 cellular genome (Table S8, Supplementary File S2). In total 820 loci containing indels larger than 2kbp were called, with an average size of 6.6kbp and a maximum of 106kbp. Among the large indels identified, 67% are insertions while 33% are deletions, and 69% are homozygous while 31% are heterozygous. Since C666-1 was originally derived from a male sample, we checked the number of indels wrongly called as heterozygous on the sex chromosomes (Table S8), and found 6 such errors among the 30 (20%) SVs identified. In addition to indels, OMSV also identified 28 medium-size inversions, 13 large inversions, 6 translocations (2 intra-chromosomal and 4 inter-chromosomal) and 68 copy number variations (CNVs) (Supplementary Files S2,S3). A translocation between intron 1 of UBR5 and intron 6 of ZNF423 was previously reported, leading to a fusion transcript (Chung et al., 2013). We were able to confirm the existence of this translocation on the list of complex SVs identified by OMSV (Figure 4).

**Figure 4:**
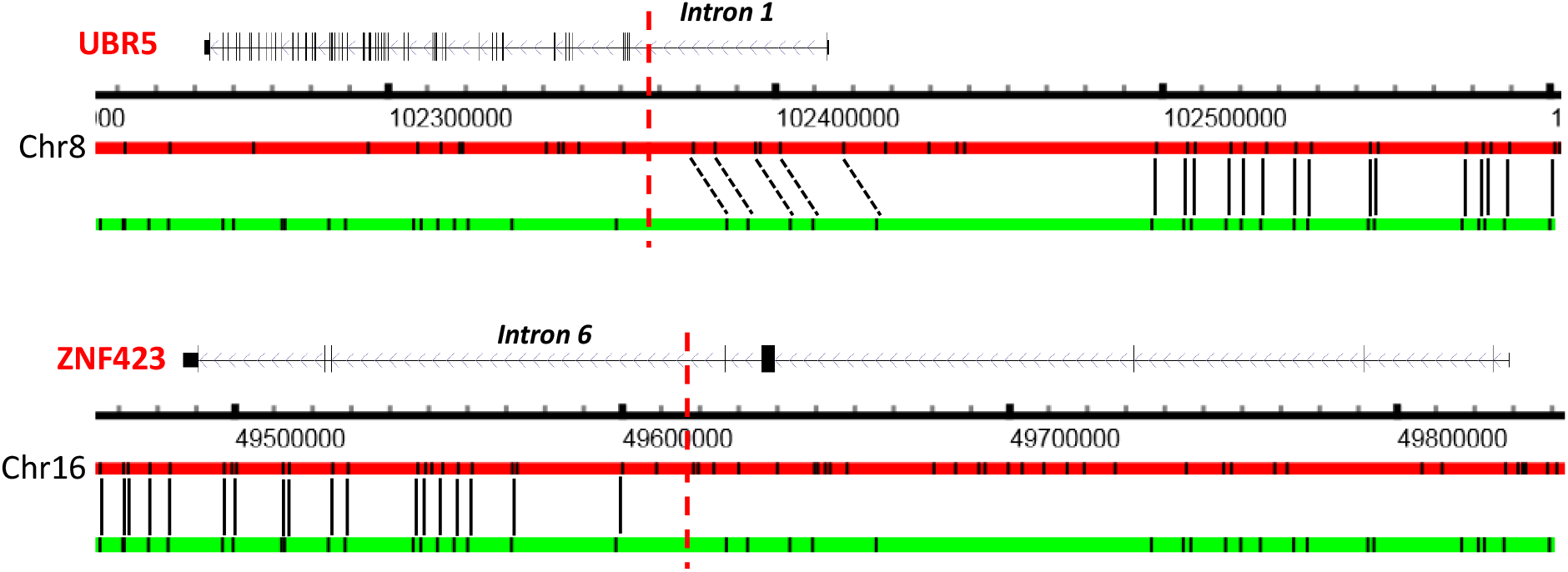
The previously reported UBR5-ZNF423 translocation in C666-1 re-identified by OMSV, visualized using the alignment view of OMView. For each of the two gene loci, the top horizontal bar shows the reference and the bottom horizontal bar shows a representative optical map. Black solid and dashed lines linking the reference and the optical map respectively represent aligned nicking sites and nicking sites that should probably be aligned but missed by the alignment pipeline. The vertical red dashed lines show the break points previously reported in Chung et al. (2013).

Whole-genome sequencing data of C666-1 were previously produced at 75x coverage with 100bp paired-end reads with an average insert size of 290bp (Tso et al., 2013). We next used two sequencing-based SV callers, Manta (Chen et al., 2016) and Pindel (Ye et al., 2009), to identify large (>2kbp) SVs from the sequencing data. Among the 820 indels identified by OMSV, 535 of them (65%) were missed by both short-read based SV callers (Figure 5a). In particular, among the 550 insertions, 439 of them (80%) were missed by both. (Figure 5a). Even for the insertions detected by Manta or Pindel, they could only provide the locations of the break points but not the sizes of the insertions, which are reported by OMSV.

**Figure 5:**
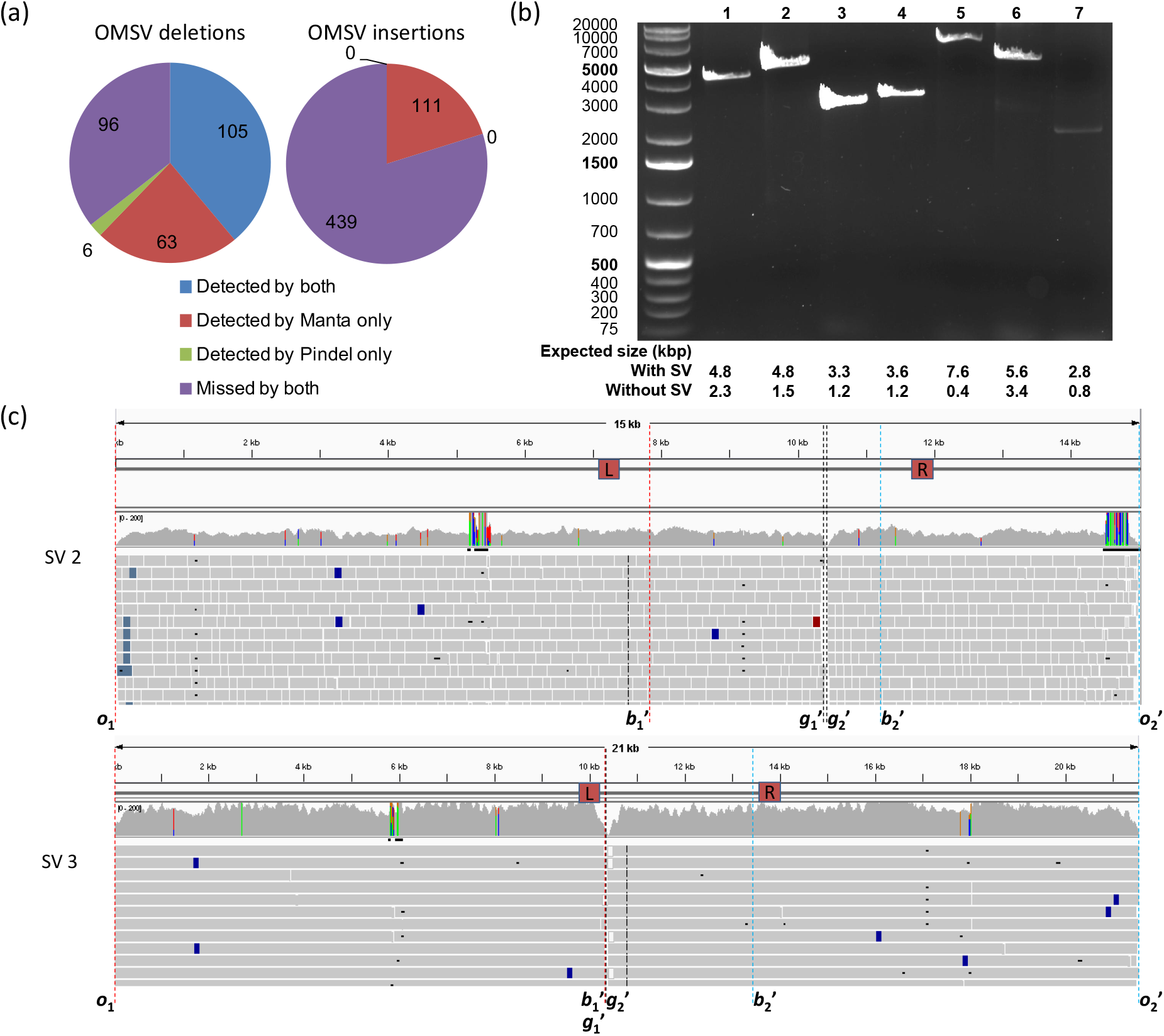
SVs identified by OMSV from C666-1. (a) Large (>2kbp) indels identified by OMSV that were also identified by one or both of the short-read based callers, Manta and Pindel, or missed by both of them. (b) PCR results of the seven selected insertions. (c) Alignment of sequencing reads to the inferred C666-1 sequences of SV 2 and SV 3. The L and R boxes mark the primer locations. Definitions of *o*_1_, 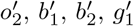 and 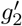 are given in Figure S9. Sequencing read alignments are visualized by IGV.

To check the accuracy of the indels identified by OMSV, we focused on insertions, which are more difficult for short-read based callers to identify accurately. Considering the maximum possible product size of PCR, we selected seven insertions for validation experiments. For each one of them, we designed primers outside its predicted break points on the reference sequence, and compared the length of the resulting PCR-amplified product with its expected length with or without the insertion (Tables S9,S10, Materials and Methods). From the PCR results (Figure 5b), all seven products confirmed the existence of the insertions.

Since optical maps only estimate SV break points up to the closest nicking sites, we used the sequencing reads to determine the break points more precisely and deduce the inserted sequences in insertions by local sequence assembly (Figure S9, Materials and Methods). The inferred sequences for the seven PCR-validated insertions and the precise SV break points are all supported by a large number of aligned sequencing reads (Figures 5c,S10).

## Discussion

In this paper, we described the OMSV pipeline for identifying SVs from nanochannel-based optical maps. The accuracy of OMSV has been confirmed by both simulations and optical maps from a family trio. OMSV outperformed the only publicly available tool for SV detection from OM data in three aspects, namely 1) OMSV identified many more SVs at a precision level similar to this method, 2) OMSV identified many of the complex SVs but this method missed all of them, and 3) OMSV ran much faster by not requiring a time-consuming de novo assembly of the optical maps.

We also used OMSV to identify SVs from the C666-1 cell line, and found 65% of them missed by sequencing-based SV callers, including 80% of the insertions. Some of these SVs were experimentally validated independently.

Currently, it is difficult to detect these SVs or estimate their sizes from sequencing data alone due to short read length and limited insert size between read pairs. Using nanochannel-based optical maps, whole SVs are easily contained in a single optical map, making SV detection highly feasible and accurate. Here we demonstrated that OMSV is a powerful tool for identifying large SVs from kilobases to more than a hundred kilobases. In fact, as long as an optical map can be correctly aligned to the reference by having sufficient nicking sites in the flanking non-SV portions, the larger an SV is, the easier for it to be detected by OMSV, since the corresponding distance change between the defining nicking sites is less likely due to scaling and measurement errors alone. This property makes OMSV an ideal complement to sequencing-based SV callers, which are generally more accurate in detecting smaller SVs.

For complex regions and very large SVs, OMSV detects them by employing a two-round alignment strategy that allows split-alignment of an optical map to multiple locations on the genome. Split-alignments of optical maps could come with a cost of extra alignment time. One way to tackle this problem is to first quickly align optical maps that can be aligned to single genomic loci using a standard aligner, and then apply the split-alignment strategy only to the remaining unaligned optical maps.

Since the SV calling modules only require a list of optical map alignments as input, the alignment methods used in the OMSV pipeline can be flexibly changed to other choices. Besides, if a high-quality *de novo* assembly of the optical maps is available, the optical maps can also be first aligned to the assembly, and their alignment to the reference can then be inferred from the further aligning the assembly to the reference. For optical maps that deviate significantly from the reference map, this two-step alignment strategy could be more accurate than directly aligning optical maps to the reference.

With each optical map coming from one DNA molecule, OMSV can potentially be extended to study haplotypes, cell type composition in a sample and cell-to-cell variability. These analyses would require highly accurate alignments of individual labels of the optical maps. Probing the nicking sites of a second enzyme using an additional color channel may further improve alignment accuracy necessary for these analyses.

We provide OMSV as open-source software, which can be used routinely in genome projects to accurately and comprehensively identify large SVs that will likely have important implications for understanding genetic diversity and disease susceptibility.

## Materials and Methods

### A complete error model for optical maps

We modeled the generation of optical maps from a DNA sequence as a random process with various types of error, which combines some ideas previously proposed (Tong et al., 2007; Tong, 2010) and several new components based on properties observed in real human optical maps (Mak et al., 2016).

In our model, the starting locations of *n* DNA fragment molecules are first uniformly and independently sampled from the DNA sequence. Each of these starting locations is used to produce a molecule with length *l*_0_ + *l_v_*, where *l*_0_ is a constant minimum molecule length, and *l_v_* is a random variable that follows a Poisson distribution with mean *μ_l_*. In real experiments, *l*_0_ is a threshold chosen such that molecules shorter than it are excluded from the analyses.

The restriction sites on each molecule can be identified by matching its sequence against the recognition motif of the nicking enzyme selected. In our model, each restriction site has a false negative rate of *f*_ for not having a corresponding observable label in the optical map due to incomplete enzymatic digestion or a measurement error.

False positive labels not originated from actual restriction sites but caused by artifacts such as nonspecific enzymatic cuts are then introduced. For every two adjacent restriction sites, the number of false positive labels is randomly sampled from a Poisson distribution with mean *df*_+_, where *d* is the distance between the two sites and *f*_+_ is the false positive rate. If the resulting number of false positive labels is non-zero, the occurrence locations of these false positives are uniformly and independently sampled from the locations between the two sites.

After these steps, each random molecule is represented by a list of distances between adjacent observed labels (including both true positives and false positives). For the convenience of discussion, we also assume the beginning and end of each molecule are marked by two artificial labels, the locations of which in actual optical maps can be determined by the span of the stained DNA backbone. Each molecule then undergoes a random stretch/compression to model sizing errors in the experiments, by multiplying the distance between every two observed labels by a factor *α*, where *α* is sampled from a Cauchy distribution with the values of the location and scale parameters set to *o_α_* and *s_α_*, respectively. We chose the Cauchy distribution since it had a good fit with the real data we produced (Figure S11).

To model the finite resolution of optical measurements, for any two adjacent labels on a stretched/compressed molecule at a distance of *d* bp from each other, they are merged into one single label that occurs at the mid-point of them with a probability of 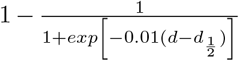 where 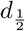 is a reference distance at which the chance for two labels to be merged is 1/2.

Finally, measurement errors are modeled by moving each label by an offset that follows a uniform distribution defined on [–*e*, *e*] for a given parameter *e*.

### SV calling modules

Based on the above generative model, we developed two statistical modules for identifying SVs from optical maps. The first module looks for individual extra or missing sites on the molecules as compared to the reference sequence. Some small SVs with only a mild change of the distance between restriction sites are better detected by this method. The second module compares the distance between two restriction sites on the molecules with that on the reference genome, which can detect larger SVs not necessarily involving extra/missing restriction sites.

Both modules require an alignment of the optical maps to a reference map obtained from the *in silico* digestion of the reference sequence, where the adjacent labels are also merged by the way described in the above section. Based on the alignments, OMSV extracts three types of information as inputs to the two SV calling modules, namely 1) the expected locations of restriction sites on the reference sequence, 2) the distance (in bp) between every two adjacent observed labels on each molecule, and 3) an alignment of the labels on the molecules to the restriction sites on the reference. Every label can be aligned to zero or one restriction site on the reference, and each restriction site on the reference can be aligned to zero or one label on each molecule.

The third module uses some additional alignment and coverage information to identify complex SVs.

### Module for identifying SVs involving extra or missing restriction sites

To identify missing restriction sites on the molecules, we adopted a method originally developed for refining optical map assemblies (Valouev et al., 2006), and extended it to detect both homozygous and heterozygous genetic variants.

Suppose there are *M* molecules aligned to a region that covers a restriction site on the reference sequence, among which *m* supports the existence of the restriction site (Figure S12a). Each of the *m* supporting molecules either actually contains the site or has a false positive label. Each of the *M* – *m* non-supporting molecules either actually does not contain the site or has a false negative. We consider three hypotheses for the observed data, namely 1) the null hypothesis 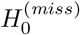 that the restriction site actually exists on the subject DNA sequence in homozygous form (and thus there are no false positives), 2) the first alternative hypothesis 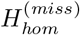 that the site is missing on the subject sequence in homozygous form (and thus there are no false negatives), and 3) the second alternative hypothesis that the site is missing on the subject sequence in heterozygous form.

Under the null hypothesis 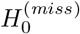, the probability of observing *m* or fewer supporting molecules is

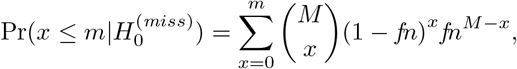

where *fn* is the false negative rate to be estimated from the observed data. Similarly, depending on whether 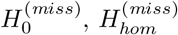 or 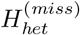 is true, the data likelihood is respectively

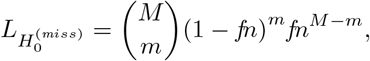

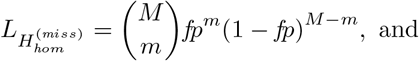

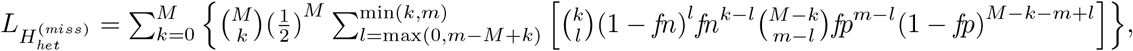

where *fp* is the false positive rate to be estimated from the observed data, *k* is, in the heterozygous case, the unknown number of molecules coming from the chromosome with the restriction site, and *l* is the number of molecules among the *k* on which the restriction site is observed. In the model, we assume there is an equal probability for a molecule to come from either chromosome.

Based on these definitions, if both the p-value 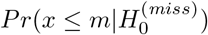 and the likelihood ratio 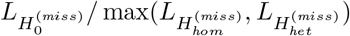 are smaller than corresponding thresholds for a site, it is considered a homozygous missing site if 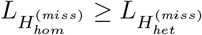 and a heterozygous missing site if 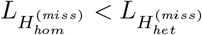.

A similar procedure is used for calling homozygous and heterozygous extra restriction sites. Suppose there are *M* molecules aligned to a region on the reference sequence, among which *m* supports the existence of a restriction site in the region that does not exist according to the reference sequence. Under the null hypothesis 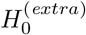 that the site is absent in homozygous form, the probability of observing m or more supporting molecules is

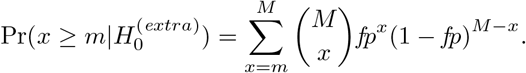

Similarly, depending on whether the site is absent in homozygous form (null hypothesis 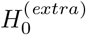), exists in homozygous form (alternative hypothesis 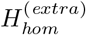), or exists in heterozygous form (alternative hypothesis 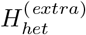), the data likelihood is respectively defined as

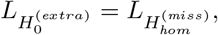

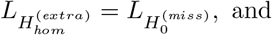

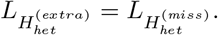

Based on these definitions, if both the p-value 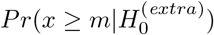 and the likelihood ratio 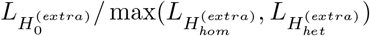 are smaller than corresponding thresholds for a site, it is considered a homozygous extra site if 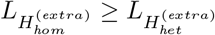 and a heterozygous extra site if 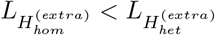.

In practice, we also define a minimum number of supporting molecules *M_min_*. For any site with less than *M_min_* molecules covering the locus (no matter supporting the presence of the restriction site or not), we did not call genetic variants from it since the result would not be reliable.

### Module for identifying SVs involving large size changes

Large SVs are usually associated with a deviation of the distance between two restriction sites on the reference sequence (Figure S12b, *d*_0_) and that on the molecules (*d*_1_), which may or may not involve extra/missing restriction sites on the molecules. To systematically identify these cases, we first check the distances between every two adjacent restriction sites on the reference sequence and compare them with the corresponding label distances on the aligned molecules (which would cover the first two cases of Figure S12b). We then check the distances between every two adjacent labels on the aligned molecules that have not been checked, and compare them with the distance between the aligned restriction sites on the reference (which would cover the third case). Each of these checks is performed by the following statistical method.

Suppose there are two (not necessarily adjacent) restriction sites on the reference sequence with a distance *d*_0_, and there are *M* aligned molecules covering the region. Suppose the distances of the corresponding aligned labels on the molecules are *d*_1_,*d*_2_,…, *d_M_*, where *d*_1_ ≤ *d*_2_ ≤ … ≤ *d_M_*. Our method computes the ratios 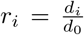 for each of the *M* molecules. It then compares the following hypotheses according to the error model we defined:

1. Null hypothesis *H*_0_, that there are no insertions or deletions between the two sites
2. *H_hom_*, that there is a homozygous indel between the two sites
3. 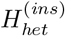, that there is a heterozygous insertion between the two sites
4. 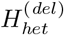, that there is a heterozygous deletion between the two sites
5. *H_tri_*, that the locus is triallelic, i.e., there are two different insertions, two different deletions, or one insertion and one deletion between the two sites, where each chromosome bears one of the two variant alleles

Under the null hypothesis *H*_0_, the likelihood of observing the distance ratios *r*_1_, *r*_2_,…, *r_M_* is

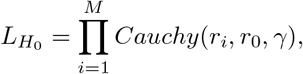

where 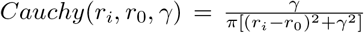 is the probability density function of the Cauchy distribution with position parameter *r*_0_ and scale parameter *γ*. In SV detection, using the Cauchy distribution to model the distance ratios has an advantage that it is not heavily affected by extreme outliers caused by alignment errors.

Under the alternative hypothesis *H_hom_*, the distance ratios *r*_1_, *r*_2_,…, *r_M_* are sampled from a Cauchy distribution with a different value for the location parameter but the same value *γ* for the scale parameter. The likelihood of observing the distance ratios is therefore

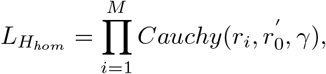

where 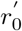 is the location parameter of the distribution of distance ratios for this indel event. Finding the maximum likelihood estimate of 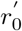 would require the use of numerical methods to solve a high-degree polynomial. Instead, we used the sample median of the *M r_i_*’s as an imperfect estimate (Hanson and Wolf, 1996).

Under the alternative hypothesis 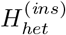, some of the distance ratios are sampled from the null distribution and the others are sampled from an alternative Cauchy distribution with a larger value 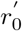 for the location parameter but the same value for the scale parameter. The likelihood of the distance ratios is 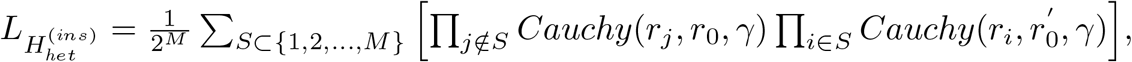 where *S* represents the set of molecules from the chromosome with the insertion, assuming an equal probability for each molecule to come from either chromosome. Practically, this likelihood is difficult to compute due to the exponential number of terms in the summation. We made an assumption that the two distributions are sufficiently separated, with 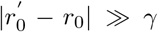. Based on this assumption, we consider only the summation terms of which *S* takes the form 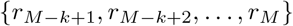, which involves only the *k* largest distance ratios. We then try all possible values of *k* such that at least *k_min_* molecules come from each chromosome (Figure S12c). As a result, the likelihood formula is simplified as 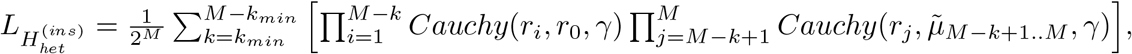 where 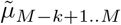 is the sample median of *r*_*M*−*k*+1_, *r*_*M*−*k*+2_,…, *r_M_*.

Similarly, for heterozygous deletions, a simplified likelihood formula is defined as 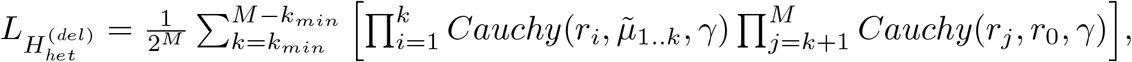, where 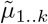, the sample median of *r*_1_, *r*_2_,…, *r_k_*, is expected to be smaller than *r*_0_ in this case (and a heterozygous deletion would not be called if this expectation is not satisfied).

For the triallelic cases, the simplified likelihood formula is defined as 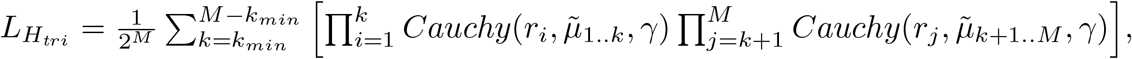, where 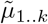 is the median of *r*_1_, *r*_2_,…, *r_k_* and 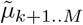 is the median of *r*_*k*+1_, *r*_*k*+2_,…, *r_M_*.

Finally, our method compares the different likelihood values. If the likelihood ratio 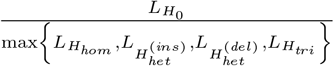 is smaller than a threshold, an SV is called according to the following rules: If max 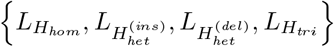 is equal to

- *L_H_hom__*: If 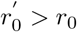, a homozygous insertion is called; Otherwise, a homozygous deletion is called.
- 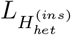: A heterozygous insertion is called.
- 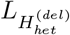: A heterozygous deletion is called.
- 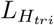: If 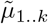 and 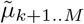 are both smaller than *r*_0_, two different deletions are called; If both are larger than *r*_0_, two different insertions are called; Otherwise, an insertion and a deletion are called.

Practically, if the distance change is too small, either absolutely or relative to the distance on the reference, the SV calls are less reliable. We therefore keep only SVs with a distance change larger than a threshold *δ*, where the distance on the molecules is defined as the median distance of the set of molecules that lead to a term with the largest value in the likelihood calculation.

### Module for identifying complex SVs

We also developed a module for identifying three types of complex SVs, namely inversions, translocations and CNVs.

#### Using split-alignment to identify large inversions and translocations

The split-alignment capability of OMBlast (Leung et al., 2017b) allows different parts of a single optical map to be separately aligned to different locations of the same chromosome (Figure S12d). The default setting of OMBlast limits the maximum distance between these different locations to reduce false alignment rate, and thus it permits direct calling of only intra-chromosomal translocations involving close loci. To detect other intra-chromosomal translocations and inter-chromosomal translocations, we used a 2-round alignment strategy (Figure S12e), in which the first round performed standard alignments of optical maps, with some optical maps only partially aligned. For these optical maps, the unaligned regions were then independently aligned again in the second round, thus allowing the detection of all types of translocations. In addition, by allowing different portions of the same optical map to be aligned in different orientations, large inversions can also be detected. To reduce false positives, only translocations and large inversions supported by two or more optical maps are considered.

#### Using reverse-palindromic CIGAR strings to identify medium-sized inversions

Inversions with size between *2kbp* and 100*kbp* can be contained in a single optical map, and are detected by locating a region in an optical map alignment with 1) a reverse-palindromic CIGAR (Compact Idiosyncratic Gapped Alignment Report) string; and 2) matching distances between adjacent restriction sites on the reference and those between adjacent labels on the reversed optical map (Figure S12f). In a CIGAR string, a matched, missing and extra label is denoted as ‘M’, ‘D’ and ‘I’, respectively. The reverse-complement of a CIGAR string is the reverse of it with ‘I’s and ‘D’s interchanged. For example, the reverse-complement of MDDI is DIIM. A CIGAR string is reverse-palindromic if it is the same as its reverse-complement, such as DIDIDI. Two distances *d*_1_ and 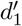 are considered matched if 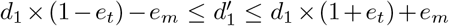, where *e_t_* and *e_m_* are the maximum scaling and measurement errors (set to 0.1 and 500bp), respectively. To control the quality, we called an inversion only if it had at least 10 supporting molecules and at least 4 nicking sites within the inverted region.

#### Using depth of coverage to identify CNVs

We modified an event-wise significance testing method (Yoon et al., 2009) to identify large CNVs. The original method uses a sliding window (with 100bp) to scan the reference and look for windows with a depth of coverage significantly different from other windows, based on the distribution of depths of windows with similar GC contents. Neighboring windows are then grouped into blocks to identify the span of the CNVs, with a method for correcting for multiple hypothesis testing. To adopt this method for OM data, first the window size was enlarged to 2*d*_1/2_ to accommodate for the lower resolution of OM data, where *d*_1/2_ is the imaging resolution. Then to determine the statistical significance of each window, instead of grouping windows by GC content, we grouped them by nicking site counts. The depths (number of aligned optical maps) of all windows within a group were fit to a Gaussian distribution, and a window was considered a CNV candidate if it received a Z-test p-value <0.05. The same procedure for determining CNV spans in the original method was then applied.

### The overall OMSV pipeline

The overall OMSV pipeline is illustrated in Figure 1b. In the alignment pipeline, we used default parameter values of RefAligner and OMBlast for all the simulated and real data except C666-1, in which case we used RefAligner parameter values for complex genomes (available on our supplementary Web site) suggested by BioNano technical team. The reference map was deduced from the human reference hg38 in all cases. RefAligner and OMBlast alignments were integrated based on the following rules:

1. If the two methods align an optical map to genomic regions within half the length of the optical map from each other, they are considered to agree on the alignment, and the alignment of RefAligner is taken.
2. If only one of the two methods can align an optical map, the alignment is taken directly.
3. If both methods cannot align an optical map, or both of them can align but their alignments do not agree with each other, the optical map is left unaligned.

We call this the “union” strategy in Figure S6. We also considered an “intersection” strategy, which only involved the alignments satisfying the first rule above.

The resulting integrated list of alignments is sent to the three modules for SV identification. The results from the three modules are then integrated to form a final list of SVs.

The parameter values of OMSV used in our experiments are listed in Table S11.

### Filtering of SVs detected from real data

We considered only optical map alignments with a confidence score of 9 or more. For the indels identified from the family trio and the C666-1 cell line, we filtered those that overlapped N-gaps, fragile sites or pseudo-autosomal regions on the reference genome. These “mask” regions are listed in Supplementary Files S4–S6. We applied the same filtering to the NA12878 SV lists obtained from sequencing-based methods. For the complex SVs, we filtered the ones located within the pseudo-autosomal regions or overlapped with the regions with ultra-high density of nicking sites.

### Generation of simulated data

We generated simulated data with either only homozygous variants or both homozygous and heterozygous variants. Two steps were involved in both cases, namely a first step for generating genomic sequences with genetic variations introduced to the human reference genome, and a second step for simulating optical maps based on the resulting genomic sequences using the error model described above.

### Simulated data with only homozygous variants

For the data set with homozygous variants only, we first downloaded the human reference sequence hg38 from the UCSC Genome Browser (Kent et al., 2002). We then generated mutations (single nucleotide variants, small and large indels, and complex SVs) on it using pIRS (profile based Illumina pair-end Reads Simulator) (Hu et al., 2012). This software was originally developed for generating short sequencing reads. We took its intermediate file containing the mutated sequence without generating the short reads. In the second step, we used the mutated sequence as input to generate simulated optical maps based on our generative model. The parameter values used in the two steps are shown in Table S12 and Table S13, respectively. The parameter values for the first step were determined based on corresponding estimates from human genomes reported in previous studies (The 1000 Genomes Project Consortium, 2010; Lu et al., 2012; Pang et al., 2010). The parameter values for the second step were estimated from our actual optical maps by aligning all molecules to the reference sequence using RefAligner, and estimating the parameter values by likelihood maximization. All these parameter values were not made known to our SV detection methods.

### Simulated data with both homozygous and heterozygous variants

For the data set with both homozygous and heterozygous variants, we generated a diploid genome as follows. It was initialized by our generated haploid genome and the reference genome as the two haplotypes. Then for each variant on the first haploid genome, it received a probability of *p_hom_* to be copied to the second haploid genome, resulting in a homozygous variant. For the remaining variants, which would remain heterozygous, each of them received a probability of *p_het_* of moving from the first haploid genome to the second. We used *p_hom_* = 0.5 and *p_het_* = 0.5 in our simulations based on a previous study (Levy et al., 2007). As a result of this procedure, the total number of SV loci in this diploid genome was the same as that in the haploid genome.

We then considered the two haploid genomes together as a diploid genome, and used the corresponding DNA sequences as the templates to produce OM data using our generative model. The parameter values used in the two steps of simulation are again shown in Table S12 and Table S13, and 26 additional data sets were generated by changing the false positive rate, false negative rate and depth of coverage, as shown in Table S1.

### Evaluation metrics of SV calling on simulated data

For the simulated data, we used the known locations of the generated SVs to compute the precision (fraction of identified SVs that are real) and recall (fraction of real SVs that are identified) rates of an SV calling method. An SV call was considered correct if it overlapped the location of a generated SV of the same type.

### Comparison with BioNano Solve

We compared OMSV with the SV caller included in BioNano Solve v3.0 (downloaded from downloadedfromhttps://bionanogenomics.com/support/software-downloads/), which was the only SV caller for nanochannel-based optical maps with publicly available software. The exact command-line arguments used can be found on the supplementary Web site.

### Evaluating the performance of OMSV in the ideal situation with no alignment errors

To estimate the performance of OMSV in the ideal situation with no alignment errors, instead of supplying optical map alignments as inputs to OMSV, we provided observed-to-reference distance ratios between neighboring nicking sites directly. For each locus, the number of distance ratios was drawn from a Gaussian distribution estimated based on the depth of coverage of the data set. The values of these distance ratios were produced by adding scaling errors to the actual distance ratio of the corresponding allele based on the sizing error parameter of the default simulated data set. The ratio of loci with and without SVs also followed the ratio in the default data set.

### Evaluation metrics of alignment pipeline on simulated data

We also defined metrics for evaluating the performance of our alignment pipeline. First, an optical map was considered correctly aligned if it was aligned to the correct haplotype of the simulated genome with the aligned location overlapping the actual location from which the optical map was generated. Alignment precision was then defined as the fraction of aligned optical maps that were correctly aligned, and recall was defined as the fraction of generated optical maps that were correctly aligned.

### Definition of Mendelian concordance

For the family trio, a locus was defined as concordant with Mendelian inheritance if the daughter’s genotype could be produced by the genotypes of the father and the mother. When zygosity was not considered, an SV identified from an individual could mean that the individual had the SV in homozygous or heterozygous form. As a result, a Mendelian error was reported only when the daughter had an SV at a locus of a type that both parents did not have. When zygosity was considered, a Mendelian error was reported when the two alleles of the daughter could not respectively come from the two parents. For this part of analysis, we considered only loci at which each of the individuals either had an SV confidently called, or it was highly unlikely that an SV could be called. The former was defined as SVs with at least 10 supporting optical maps and a likelihood ratio of at most 10^-6^ for each other hypothesis. The latter was defined as cases in which an SV could not be called even at the loose thresholds of 4 supporting optical maps and a likelihood ratio of 1.

### Computation of expected Mendelian concordance

In order to check whether the observed Mendelian concordance values of the SVs identified from the trio were consistent with the precision and recall estimates of our simulation, we computed the expected Mendelian concordance values as follows. First, we estimated the probabilities *P*(*G*_2_|*G*_1_) where *G*_1_ and *G*_2_ are respectively the actual genotype and the genotype called by OMSV, each with possible alleles *A* (the reference allele) and *a* (the alternative allele). The probabilities *P*(*aa*|*aa*), *P*(*Aa*|*aa*), *P*(*AA*|*aa*), *P*(*aa*|*Aa*), *P*(*Aa*|*Aa*) and *P*(*AA*|*Aa*) were all estimated based on the fraction of homozygous and heterozygous variants generated in our simulated data that were called by OMSV to have the corresponding genotypes. For the remaining three conditional probabilities, *P*(*Aa*|*AA*) = *P*(*AA*|*Aa*)*P*(*Aa*)/*P*(*AA*) ≈ *P*(*AA*|*Aa*)*P*(*Aa*), where *P*(*AA*|*Aa*) was again estimated from our simulation result, and *P*(*Aa*) was estimated as half the prior SV probability of the human genome, 8 × 10^-3^/2 (based on the median total SV size of 20Mbp per individual reported in Sudmant et al. (2015)), assuming equal probability for homozygous and heterozygous SVs. *P*(*aa*|*AA*) was estimated in exactly the same way. Finally, *P*(*AA*|*AA*) = 1 — *P*(*Aa*|*AA*) — *P*(*aa*|*AA*).

With all these 9 probabilities computed, we charted the probability for each combination of actual and called genotypes of the trio. Specifically, the father, mother and daughter genotypes were denoted as a triple. For example, (*AA*, *aa*, *Aa*) represents the situation that the father has the reference genotype, the mother has an SV in homozygous form and the daughter has the SV in heterozygous form. The probability for an actual genotype combination *C*_1_ to be called as a genotype combination *C*_2_ was calculated as the product of the three corresponding conditional probabilities, assuming SV calling errors of the three individuals are independent. For example, *P*((*AA*, *AA*, *aa*)|(*AA*, *aa*, *Aa*)) = *P*(*AA*|*AA*)*P*(*AA*|*aa*)*P*(*aa*|*Aa*).

When zygosity was considered, the actual genotype combination must come from the set of 15 combinations concordant with Mendelian inheritance, *O* = {(*AA*, *AA*, *AA*), (*AA*, *Aa*, *AA*), (*AA*, *Aa*, *Aa*), (*AA*, *aa*, *Aa*), (*Aa*, *AA*, *AA*), (*Aa*, *AA*, *Aa*), (*Aa*, *Aa*, *AA*), (*Aa*, *Aa*, *Aa*), (*Aa*, *Aa*, *aa*), (*Aa*, *aa*, *Aa*), (*Aa*, *aa*, *aa*), (*aa*, *AA*, *Aa*), (*aa*, *Aa*, *Aa*), (*aa*, *Aa*, *aa*), (*aa*, *aa*, *aa*)}. The overall expected Mendelian concordance rate was then calculated as 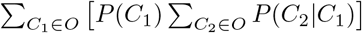. We estimated the prior probabilities *P*(*C*_1_) by the number of times such genotype combination was called by OMSV in the trio data.

When zygosity was ignored, the expected Mendelian concordance rate was calculated as 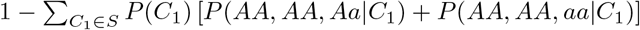.

### Comparing with sequencing-based results for NA12878 SVs

We lifted over the SV lists of NA12878 from Parikh et al. (2016) and Sudmant et al. (2015) from hg19 to hg38. We then filtered both these lists and our list of SVs by removing SVs with a size smaller than 2,000bp or overlapping the mask regions. The remaining SVs on the three lists were then compared.

## Production of optical maps from C666-1

### High-molecular-weight DNA extraction

The cancer cell line was washed with PBS and spun down to pellet. 10^6^ cell/mL were obtained upon resuspension in PBS, and embedded in 1.5% low-melting agarose plugs in 0.5x TBE (CHEF Genomic DNA Plug Kit, Bio-Rad). Subsequent handling of the DNA followed BioNano Genomics recommended protocols: the agarose plugs were incubated with proteinase K with Lysis Buffer at 50°C overnight. The plugs were washed by Wash Buffer to stabilize DNA in plugs, and the quality was assessed using pulsed-field gel electrophoresis. A plug was then washed with TE buffer and melted in 70°C. After being solubilized with 0.4 U of GELase (Epicentre), the purified DNA was subjected to 2.5hr of drop-dialysis and was shredded by 9 strokes of gentle pipetting. The viscous DNA was allowed to equilibrate overnight at room temperature to increase homogeneity. It was then quantified using a Qubit Broad Range dsDNA Assay Kit (Life Technologies).

### DNA labeling

The DNA was labeled using the IrysPrep Reagent Kit (BioNano Genomics). Specifically, 300 ng of purified genomic DNA was nicked with 0.3U of nicking endonuclease Nt.BspQI (New England BioLabs) at 37° for 2 hr in buffers BNG3. The nicked DNA was labeled with a fluorescent-dUTP nucleotide analog using Taq polymerase (NEB) for 1 hr at 72°. After labeling, the nicks were ligated with Taq ligase (NEB) in the presence of dNTPs. The backbone of fluorescently labeled DNA was counterstained with YOYO-1 (BioNano Genomics IrysPrep Reagent Kit).

### Data collection and assembly

The DNA was loaded onto the BioNano Genomics IrysChip and linearized and visualized by the Irys system. The DNA backbone length and locations of fluorescent labels along each molecule were detected using the Irys software. Single-molecule maps were assembled *de novo* into genome maps using the IrysSolve software tools developed at BioNano Genomics (Cao et al., 2014).

### Identifying SVs from C666-1 using short reads

We used the default settings of Manta and Pindel to identify SVs from the sequencing data of C666-1. We considered only large (>2kbp) SVs supported by at least 20 reads/read pairs.

### Selection of C666-1 SVs for experimental validations

We selected SVs identified by OMSV from C666-1 cells for experimental validations based on the following criteria: 1) We only selected insertions, since large insertions are particularly difficult to identify and their sizes difficult to determine from sequencing reads alone, 2) We selected SVs of predicted sizes smaller than 10kbp, since sequences longer than 10kbp can hardly be amplified by PCR, 3) We only selected SVs determined by OMSV as homozygous, such that validation results can be more easily interpreted. We designed PCR primers from regions outside the predicted SV break points on the reference genome sequence while avoiding repetitive sequences (Tables S9, S10).

### Integrating sequencing reads to infer precise break points and inserted sequences

For each SV identified by OMSV from C666-1 that occurs within the region [*o*_1_, *o*_2_] of the human reference genome sequence hg38 with an estimated size of *s*, we performed the following steps (Figure S9, Table S9):

1. Construct a tentative C666-1 sequence by replacing the region [*o*_1_, *o*_2_] by *x* copies of *N* (i.e., unknown) nucleotides, where *x* = *o*_2_ – *o*_1_ + *s* for an insertion and *x* = *o*_2_ – *o*_1_ – *s* for a deletion.
2. Use GapCloser (Luo et al., 2012) to infer the actual sequence of this *N* region based on local assembly of sequencing reads and the flanking sequences, which may or may not resolve all the *N*s.
3. Align sequencing reads to the region [*o*_1_, *o*_2_] of the reference sequence using BWA (Li and Durbin, 2009), visualizing only read pairs with both sides aligned using IGV (Robinson et al., 2011).
4. Align sequencing reads to the inferred C666-1 sequence using BWA, visualizing only read pairs with both sides aligned.
5. Use the alignment results to evaluate confidence of the SV, the break points and the inserted sequences in the case of insertions.

## Acknowledgments

T-FC and KYY are partially supported by the HKSAR Food and Health Bureau Health and Medical Research Fund HMRF12110542. KWL, T-FC and KYY are partially supported by the HKSAR Research Grants Council Theme-based Research Scheme T12-401/13-R.

